# Mitochondrial defects and metabolic vulnerabilities in Lynch syndrome-associated MSH2-deficient endometrial cancer

**DOI:** 10.1101/2024.06.10.596841

**Authors:** Mikayla Borthwick Bowen, Brenda Melendez, Qian Zhang, Diana Moreno, Leah Peralta, Wai Kin Chan, Collene Jeter, Lin Tan, M. Anna Zal, Philip L. Lorenzi, Kenneth Dunner, Richard K Yang, Russell R. Broaddus, Joseph Celestino, Nisha Gokul, Elizabeth Whitley, Rosemarie Schmandt, Karen Lu, Hyun-Eui Kim, Melinda S. Yates

## Abstract

Lynch syndrome (LS) is defined by inherited mutations in DNA mismatch repair genes, including *MSH2,* and carries 60% lifetime risk of developing endometrial cancer (EC). Beyond hypermutability, specific mechanisms for LS-associated endometrial carcinogenesis are not well understood. Here, we assessed the effects of MSH2 loss on EC pathogenesis using a novel mouse model (PR-Cre *Msh2*^flox/flox^, abbreviated Msh2KO), primary cell lines established from this model, human tissues, and human EC cell lines with isogenic MSH2 knockdown. Beginning at eight months of age, 30% of Msh2KO mice exhibited endometrial atypical hyperplasia (AH), a precancerous lesion. At 12 to 16 months of age, 47% of Msh2KO mice exhibited either AH or ECs with histologic features similar to human LS-related ECs. Transcriptomic profiling of EC from Msh2KO mice revealed a transcriptomic signature for mitochondrial dysfunction. Studies *in vitro* and *in vivo* revealed mitochondrial dysfunction based upon two mechanisms: marked mitochondrial content reduction, along with pronounced disruptions to the integrity of retained mitochondria. Human LS-related ECs also exhibited mitochondrial content reduction compared with non-LS-related ECs. Functional studies revealed metabolic reprogramming of MSH2-deficient EC cells *in vitro*, including reduced oxidative phosphorylation and increased susceptibility to glycolysis suppression. We are the first to identify mitochondrial dysfunction and metabolic disruption as a consequence of MSH2 deficiency-related EC. Mitochondrial and metabolic aberrations should be evaluated as novel biomarkers for endometrial carcinogenesis or risk stratification and could serve as targets for cancer interception in women with LS.

**Significance:** This is the first study to report mitochondrial dysfunction contributing to MSH2-deficient endometrial cancer development, identifying a noncanonical pathway for MSH2 deficient carcinogenesis, which also imparts vulnerability to metabolic targeting.

## 1. Introduction

Lynch syndrome (LS) is the most common cancer susceptibility syndrome, affecting 1 in 279 individuals in the United States (1). LS increases risk of multiple cancer types, most commonly endometrial cancer (EC) and colorectal cancer (CRC). LS is an autosomal dominant inherited syndrome defined by a loss-of-function mutation in a DNA mismatch repair (MMR) gene, namely *MLH1*, *MSH2*, *MSH6*, or *PMS2* (2). Loss of DNA mismatch repair, termed mismatch repair deficiency (MMRd), leads to increased DNA mutations and thus increased likelihood for oncogenic mutations to occur. LS is associated with up to 60% lifetime risk of EC, and approximately 2-5% of all diagnosed ECs are attributed to LS (3,4). Major knowledge gaps remain in understanding EC pathogenesis and opportunities for improved risk stratification and cancer interception are critically needed. Prior studies in colon cancer or stem cells have shown increased susceptibility of MMRd cells to oxidative stress (5–7) and ionizing radiation (8–10). Mouse models with global loss of mismatch repair genes *Msh2*, *Pms2*, or *Mlh1* produce lymphomas at an early age (11–13), which has been a limiting factor to studying tumorigenesis in other tissues. Translational studies on LS-related EC development have been limited, in part due to lack of a mouse model for LS-related EC. We report here the first mouse model for LS-related EC development through MSH2 loss targeted to the female reproductive tract, which facilitates mechanistic studies and will revolutionize translational research in LS-EC. Elucidating mechanisms of LS-related EC development will be critical in providing novel biomarkers for risk stratification and/or therapeutic targets for prevention and interception.

Mitochondrial dysfunction has long been implicated in cancer development and progression due to its effects on metabolic reprogramming, generation of reactive oxygen species, hypoxia signaling, and apoptosis signaling (reviewed in (14)). In EC, studies have evaluated the impact of hyperestrogenism on mitochondria. Because estrogen stimulates mitochondrial biogenesis, EC is typically associated with increased mitochondrial content (15). Indeed, type I ECs, an outdated classification of ECs associated with hyperestrogenism, exhibit elevated mitochondrial biogenesis compared to benign endometrium (16). However, a subset of ECs from The Cancer Genome Atlas (TCGA) exhibits transcriptional signatures indicative of low mitochondrial content and function (17). Importantly, these “OXPHOS-low” ECs exhibited significantly increased prevalence of MMRd compared to ECs with intact OXPHOS signaling. The prior associations of mitochondrial dysfunction with MMRd colorectal cancer (18) and impaired oxidative phosphorylation signaling in MMRd ECs suggest a potential link between LS and/or MMRd and mitochondrial disruption. Our studies probe this critical link in MSH2-deficient EC using a novel mouse model.

This study reports a valuable new mouse model of LS-EC and defines a role for MSH2 loss in mitochondrial defects and metabolic reprogramming. We utilize this novel mouse model, including multiple cell lines derived from this system, in parallel with tissue from patients with LS to interrogate the importance of this phenotype in development of LS-EC. Here, we are the first to identify mitochondrial dysfunction as a mechanism for LS-EC development. Mitochondrial dysfunction and metabolic disruption are consequences of MSH2 deficiency-related EC and have important implications for cancer interception or therapeutic approaches for LS-EC. In addition, mitochondrial content loss and dysfunction should be evaluated as potential shared mechanisms for EC development through other risk conditions for EC development.

## 2. Results

### 2.1 Mouse model for MSH2 loss targeted to the reproductive tract develops endometrial pathology similar to human LS-ECs

To achieve *Msh2* loss in the endometrium, we bred progesterone receptor Cre knock-in (*PR-Cre*) mice (19) to *Msh2^LoxP/LoxP^* mice (20) to produce *PR-Cre*+*Msh2^LoxP/LoxP^* mice with targeted loss of MSH2 expression in tissues that express PR, including the uterus, ovaries, and mammary glands (**Figure 1A-B**). Female mice expressing *PR-Cre* and *Msh2^LoxP/LoxP^* (hereafter abbreviated Msh2KO) developed endometrial pathology as early as eight months of age, with 58% of Msh2KO mice exhibiting a pre-cancerous atypical hyperplasia (AH) lesion. By 12 to 16 months of age, 24% of Msh2KO mice developed EC and an additional 23% exhibited AH (**Table 1**). Control mice (*Msh2^LoxP/LoxP^* without *PR-Cre*) did not develop tumors and rarely (4%) developed AH. Msh2KO tumors ranged from microscopic lesions to large tumors involving an entire uterine horn. Tumors that developed in Msh2KO mice exhibited histology similar to human endometrioid (50%), serous (40%), or mixed endometrioid with serous and/or mucinous (10%) subtypes and all were microsatellite instable. Representative images of mouse tissues throughout the Msh2KO EC development spectrum are shown in **Figure 1C**.

**Figure 1.**
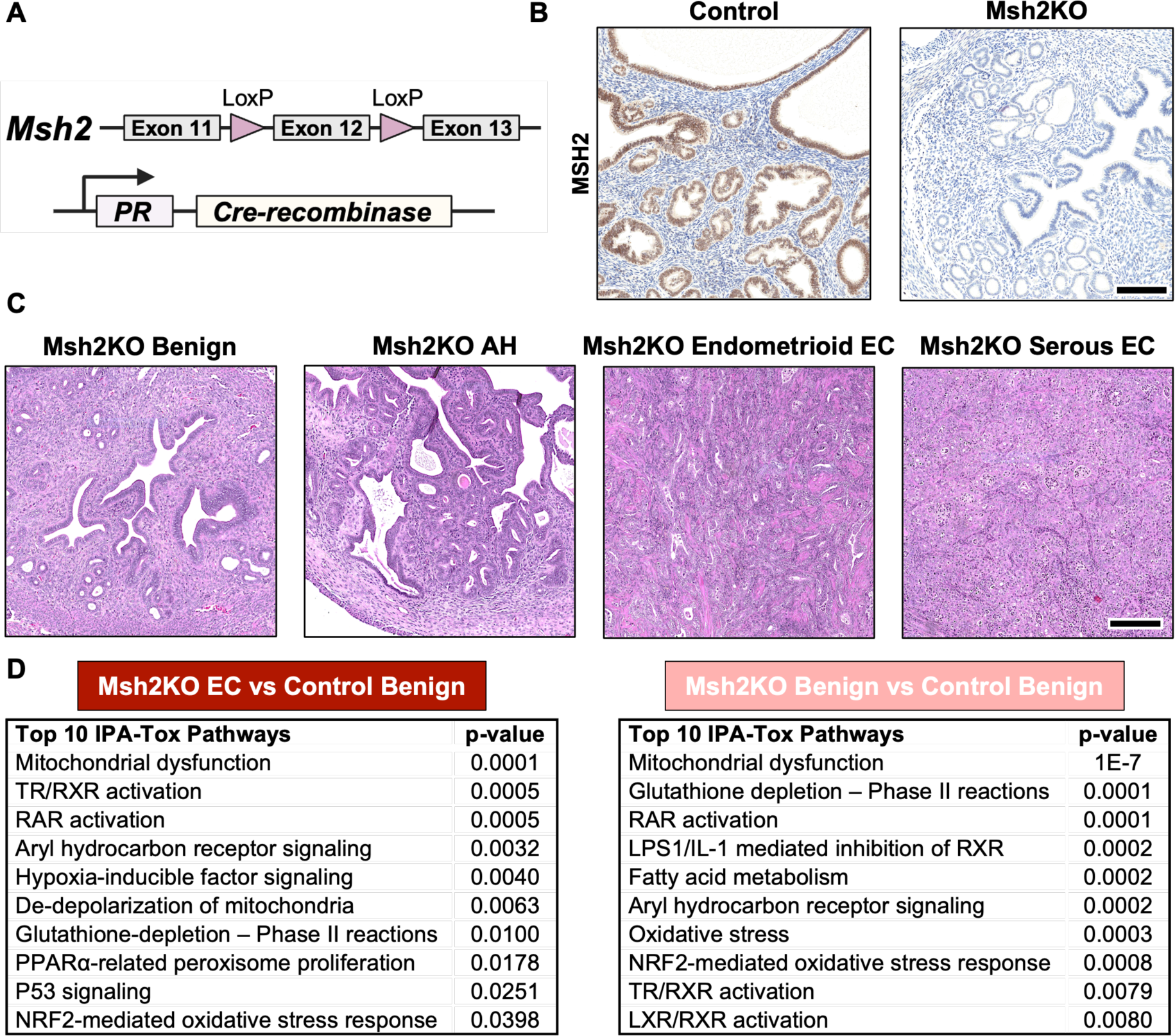
Characterization of mouse model for MSH2-deficient EC development. **(A)** Msh2KO mice express PR-Cre and a floxed Msh2 gene segment. **(B)** Presence of MSH2 by immunohistochemistry (IHC) in control mouse endometrium (left) and Msh2KO mouse endometrium (right). Scale bar=200 μm. **(C)** Representative images of hematoxylin and eosin (H&E) stained uterine tissues from Msh2KO mice with benign endometrium, Msh2KO mice with AH, and Msh2KO mice with endometrioid EC and serous EC. Scale bar=200 μm. **(E)** Top ten deregulated pathways in Msh2KO EC (left) or Msh2KO benign endometrium (right) versus control benign endometrium identified by Ingenuity Pathway Analysis (IPA) using differentially expressed genes from ClariomD microarray transcriptome analyses.

**Table 1.**
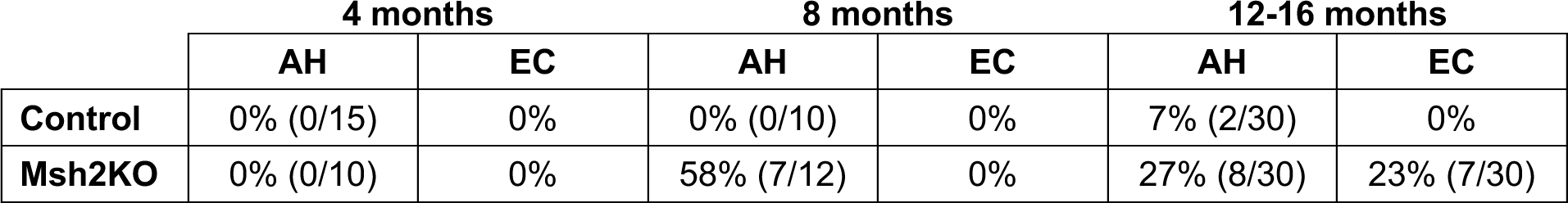
Incidence of endometrial pathologies in Control versus Msh2KO mice.

|  | 4 months |  | 8 months |  | 12-16 months |  |
| --- | --- | --- | --- | --- | --- | --- |
|  | AH | EC | AH | EC | AH | EC |
| **Control** | 0% (0/15) | 0% | 0% (0/10) | 0% | 7% (2/30) | 0% |
| **Msh2KO** | 0% (0/10) | 0% | 58% (7/12) | 0% | 27% (8/30) | 23% (7/30) |

### 2.2 MSH2 loss revealed transcriptional signature for mitochondrial dysfunction in Msh2KO mice

We profiled an initial cohort of Msh2KO tumors (n = 7) and Msh2KO mice with benign endometrium (n = 6) compared to age-matched control mice (n = 4) using transcriptomic profiling of microdissected glands via microarray analysis (Clariom D, Affymetrix). A total of 621 genes were differentially expressed between control and Msh2KO benign endometrium with fold change values of > 2 or < -2 and false discovery rate (FDR) <0.05. Between control benign endometrium and Msh2KO EC, 2,083 genes were differentially expressed with fold change values of > 2 or < -2 and FDR<0.05. Pathway analysis (Ingenuity) of differentially expressed genes revealed 88 dysregulated pathways in Msh2KO benign endometrium and 101 dysregulated pathways in Msh2KO EC compared to control benign endometrium (p<0.05) (**Figure 1D**). The top pathway significantly differing in both Msh2KO benign endometrium and EC compared to control benign endometrium was “Mitochondrial dysfunction,” where the overall panel of mitochondrial genes in the pathway exhibited 0.3-fold (Msh2KO benign) and 0.2-fold (Msh2KO EC) expression compared to control benign endometrium.

### 2.3 Msh2KO EC exhibits loss of mitochondrial content

With transcriptomic signatures pointing to mitochondrial dysfunction, including reduced expression of multiple mitochondrial electron transport chain complex genes in MSH2-deficient EC development, we evaluated overall mitochondrial content at the protein level. Immunohistochemistry (IHC) for the mitochondrial surface marker TOM20 (translocase of the outer mitochondrial membrane complex subunit 20) throughout the EC development spectrum showed that Msh2KO endometrial tumors exhibited significantly reduced TOM20 staining (**Figure 2**). Percent of cells with low TOM20 expression (TOM20-low), indicating low mitochondrial content, were increased in Msh2KO tumor tissue (30%) compared to control (16.3%) (p<0.01). Percent of cells with high TOM20 expression (TOM20-high), indicating high mitochondrial content, was decreased in Msh2KO tumor tissue (4.78%) compared to control endometrium (14.29%) (p<0.01) (**Figure 2B)**. Additionally, percent of TOM20-high cells was reduced in the Msh2KO AH tissue (4.43%, p<0.01), indicating that mitochondrial content reduction may begin at the precancerous AH stage. Zoomed insets (40X) show staining pattern differences: whereas control benign endometrium has strong staining throughout the cytoplasm, Msh2KO benign endometrium and AH tissues show patchy areas of low staining. Msh2KO EC shows low staining with focal perinuclear positivity.

**Figure 2.**
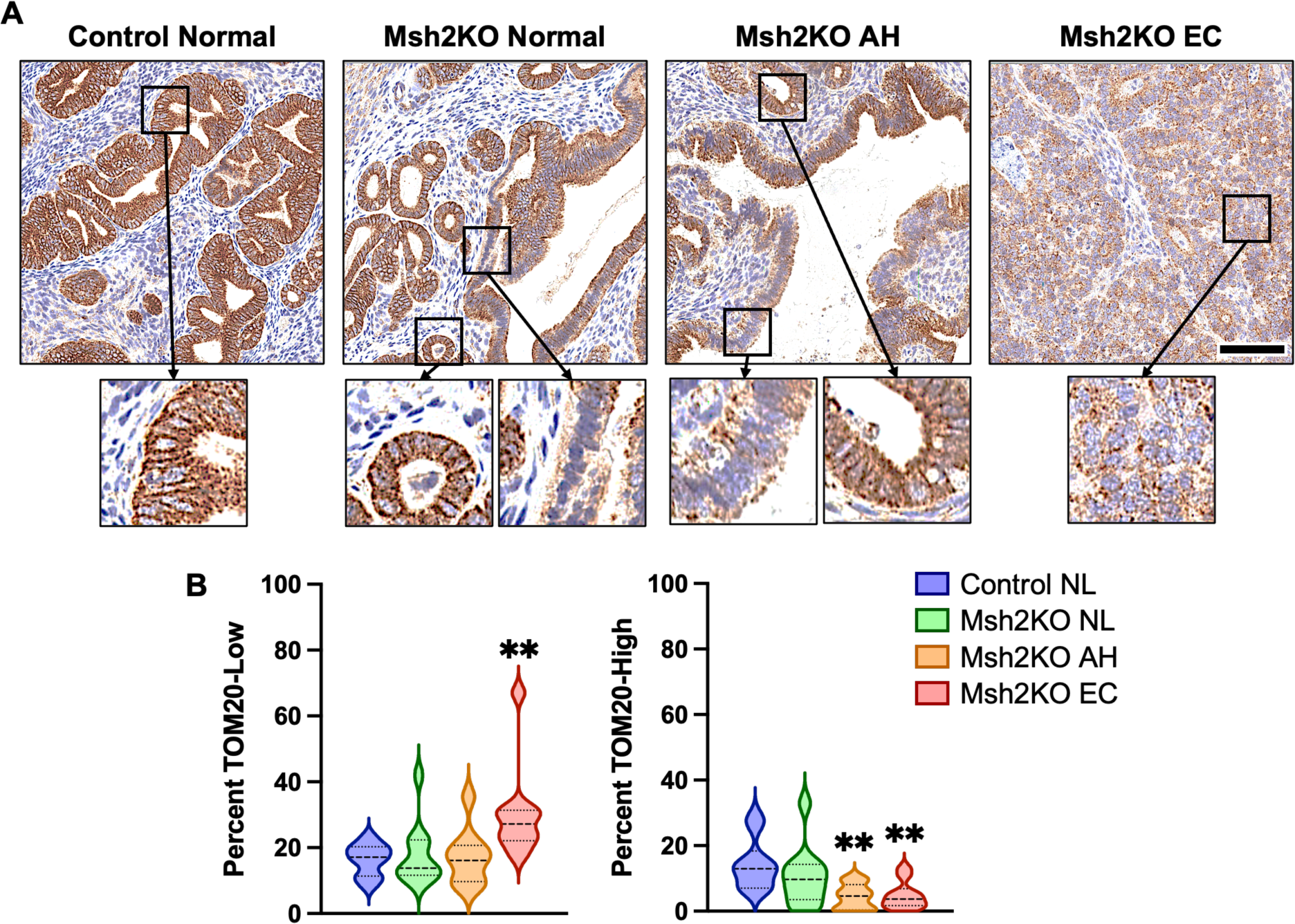
Mitochondrial content decreased during Msh2KO EC development. **(A)** Representative images (20X) of TOM20-stained control normal endometrium, Msh2KO normal endometrium, Msh2KO AH, and Msh2KO EC IHC specimens. Scale bar=100 μm. Insets 40X. Staining of glandular areas was quantified using ImageScope Cytoplasmic v2 algorithm. **(B)** Percent cells with low (left) and high (right) TOM20 staining. **p<0.01 compared to control normal (NL) endometrium. Comparisons made using ANOVA with Dunnett’s multiple comparisons test.

To evaluate the relevance to human LS-EC and determine whether mitochondrial content loss is specific to MSH2-deficient EC rather than reflective of carcinogenesis more broadly, we conducted IHC staining against TOM20 in non-MMRd human ECs and those with MSH2-deficiency (**Figure 3A**). Percent TOM20-high cells were reduced by 20.2% and percent TOM20-low cells were increased by 20.2% in MSH2-deficient ECs compared to non-MMRd ECs (**Figure 3B**), further supporting that mitochondrial content reduction is an MSH2 deficiency-related consequence in EC.

**Figure 3.**
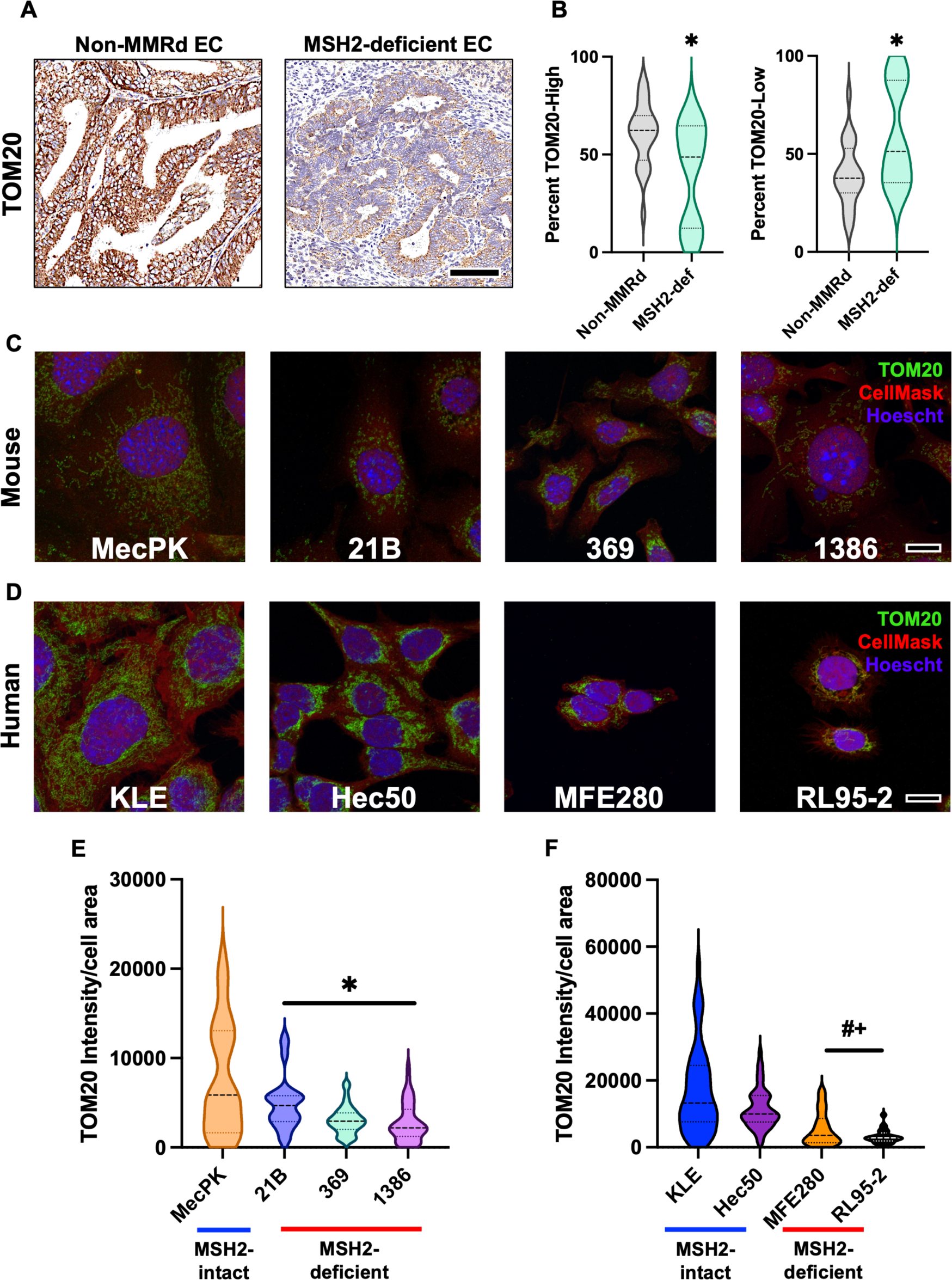
MSH2-deficient EC exhibited mitochondrial content reduction. **(A)** Representative images of TOM20-stained human non-MMRd EC (left) and MSH2-deficient EC (right) by IHC. Scale bar = 100μm. Staining was quantified using ImageScope Cytoplasmic algorithm. **(B)** Percent cells with high (left) and low (right) TOM20 staining. *p<0.05 compared to non-MMRd. Comparisons made using Mann-Whitney U test. **(C)** Mouse and human **(D)** EC cell lines were stained using immunofluorescence with TOM20 (green) and HCS CellMask (red) and counterstained with Hoescht 33342. Representative z-stacked Airyscan-processed images taken using a confocal microscope and Airyscan detector shown. Scale bar=10 μm. Mitochondrial content (TOM20 intensity) corrected for cell area (HCS CellMask) was quantified using CellProfiler for mouse **(E)** and human **(F)** EC cell lines. *p<0.01 relative to MecPK. ^#^p<0.0001 relative to KLE. ^+^p<0.05 relative to Hec50. Comparisons made using ANOVA with Dunnett’s multiple comparisons test.

A panel of MSH2-deficient mouse and human cell lines were evaluated using immunofluorescent imaging (IF) of TOM20-stained cells. These *in vitro* studies confirmed that MSH2-deficient mouse and human cells similarly exhibited reductions in mitochondrial content (TOM20 intensity relative to cell size [HCS CellMask]) compared to MSH2-intact counterparts (**Figure 3D-G**). Compared to MSH2-intact mouse EC cells (MecPK), the unique EC cell lines derived from three different Msh2KO tumors (21B, 369, and 1386) exhibited 36%, 60%, and 64% decreases in TOM20 intensity, respectively (p<0.01). Compared to MSH2-intact human EC cells (KLE), MSH2-deficient MFE280 and RL95-2 cells had 68% and 81% decreases in TOM20 intensity (p<0.0001), signifying that MSH2-deficient mouse and human EC cell lines exhibit mitochondrial content reduction. An additional MSH2-intact human EC cell line (Hec50) had reduced TOM20 intensity compared to KLE cells (by 33%) but had greater TOM20 intensity than both MFE280 and RL95-2 (by 52% and 72%, respectively) (p<0.05).

### 2.4 MSH2-deficient endometrial cancer cells exhibit defects in mitochondrial integrity and function

We next evaluated the consequences of MSH2 loss on specific alterations in mitochondrial integrity and function. Transmission electron microscopy (TEM) of mitochondria in MSH2-intact and -deficient EC cell lines revealed abnormalities in mitochondrial ultrastructure, including cristae malformations and mitochondrial condensation (darkened mitochondrial matrix) and swelling (enlarged mitochondrial matrix with malformed cristae) (**Figure 4A**). Because these ultrastructural changes are indicative of disrupted membrane polarization, we performed flow cytometric analyses of JC-1-stained cells to measure mitochondrial membrane potential (MMP) in MSH2-deficient and -intact cells. Both mouse and human MSH2-deficient EC cells exhibited reductions in MMP compared to MSH2-intact counterparts (**Figure 4B-C**). Furthermore, isogenic MSH2-knockdown decreased MMP in KLE and Hec50 cells (**Figure 4D-F**), though not in the KLE-shRNA2 line.

**Figure 4.**
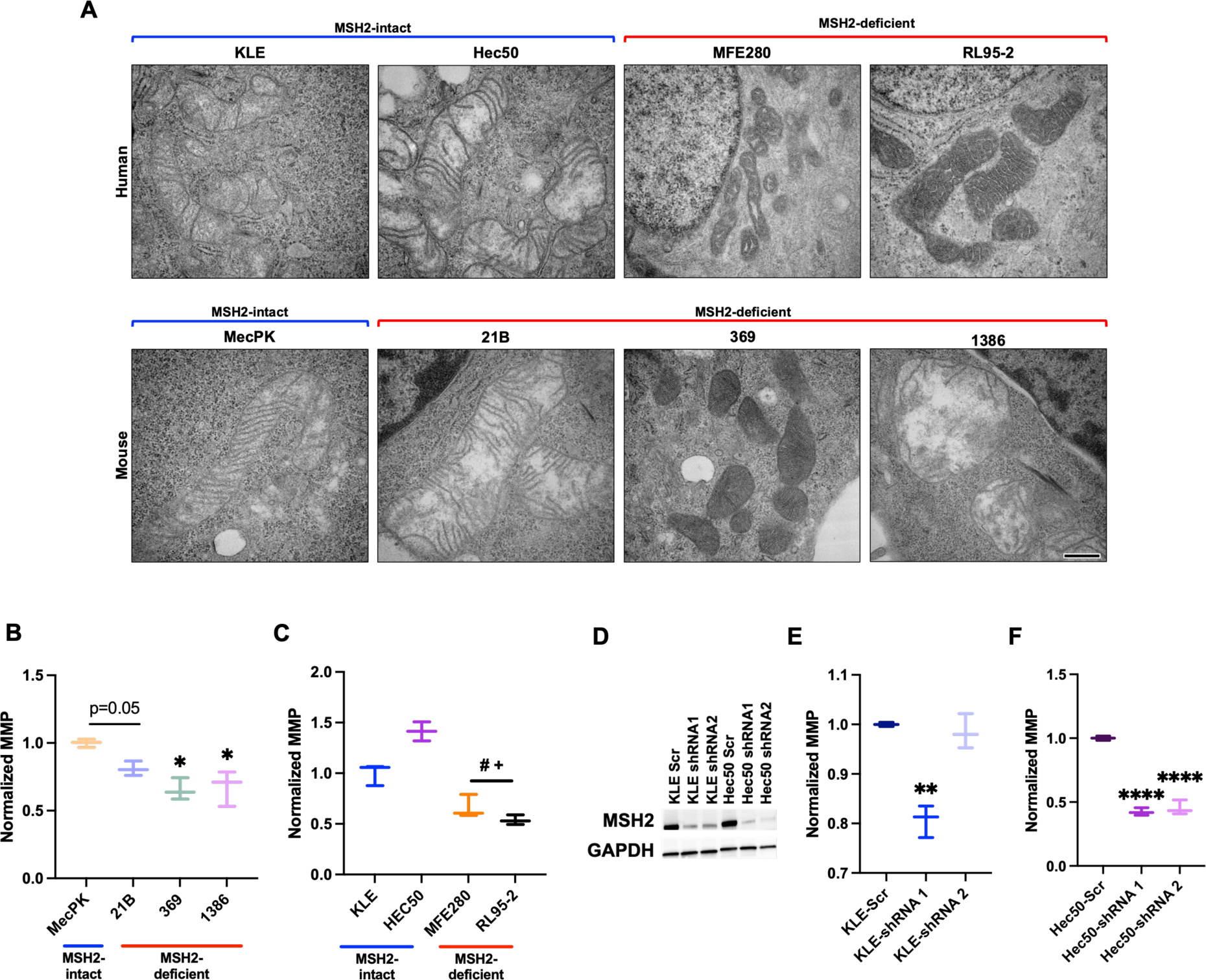
MSH2-deficient EC cells exhibit defects to mitochondrial integrity. **(A)** Transmission electron microscopy was performed on human (top row) and mouse (bottom row) MSH2-intact and MSH2-deficient EC cell lines. Scale bar = 200 nm. Mitochondrial membrane potential was measured using flow cytometric analysis of JC-1-stained mouse **(B)** and human **(C)** EC cells. *p<0.05 relative to MecPK. ^#^p<0.05 relative to KLE. ^+^p<0.05 relative to Hec50. Comparisons made using ANOVA with Dunnett’s multiple comparisons test. **(D)** Western blot of MSH2 expression (GAPDH loading control) showing lentiviral shRNA-mediated MSH2 knockdown was achieved in KLE and Hec50 cells using shRNA1 and shRNA2 sequences (see Supplemental figure _ for vector design). Mitochondrial membrane potential was measured using flow cytometric analysis of JC-1-stained KLE **(E)** and Hec50 **(F)** cells with (shRNA1, shRNA2) and without (Scr) MSH2 knockdown. **p<0.01 relative to KLE-Scr. ****p<0.0001 relative to Hec50-Scr. Comparisons made using ANOVA with Dunnett’s multiple comparisons test.

Maintenance of intact MMP is essential for mitochondrial respiration, a major metabolic function of mitochondria. To measure functional consequences of defects in mitochondrial content and membrane potential as described, we performed mitochondrial stress tests (MSTs) on MSH2-deficient and -intact mouse and human cells (**Figure 5**). MSH2-deficient mouse and human EC cells exhibited reductions in both baseline mitochondrial function (basal oxygen consumption rate [OCR]) and induced mitochondrial function (spare respiratory capacity [SRC]) compared to MSH2-intact counterparts (**Figure 5A-D**). Furthermore, MSH2 knockdown decreased both baseline and induced mitochondrial function in KLE and Hec50 cells (**Figure 5E-H**). Of note, all isogenic lines with MSH2 knockdown exhibited reductions in baseline and induced mitochondrial function, including the KLE shRNA 2 line that lacked differences in MMP above (see Figure 4). Based upon these results, mitochondrial content, membrane potential, and respiration are disrupted in MSH2-deficient EC, causing mitochondrial dysfunction.

**Figure 5.**
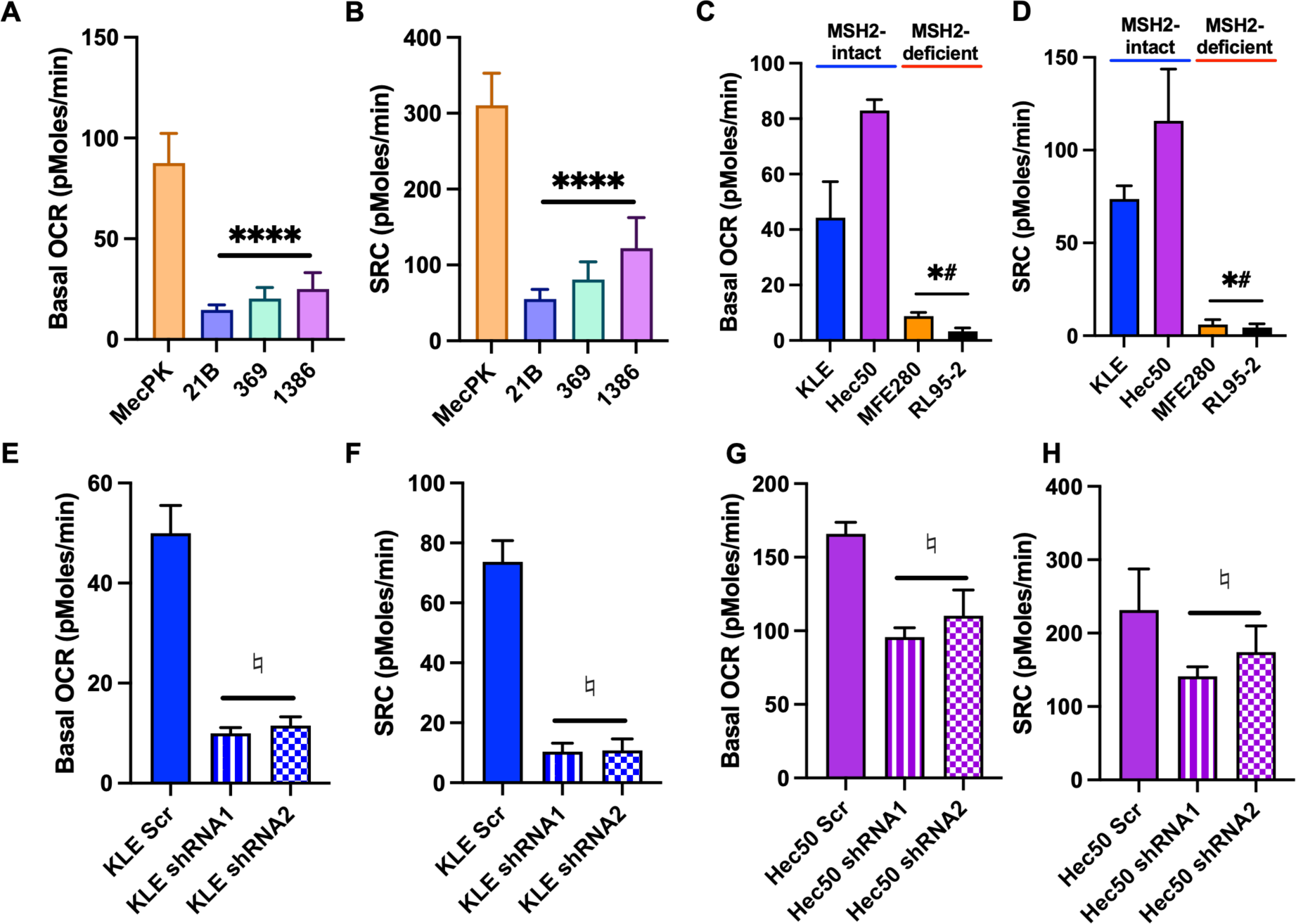
MSH2 loss confered reduced mitochondrial function in EC cells. Mitochondrial stress tests (MSTs) were performed on mouse (A-B) and human (C-D) EC cells, as well as on isogenic KLE (E-F) and Hec50 (G-H) cells with and without lentiviral shRNA-mediated MSH2 knockdown. Oxygen consumption rate (OCR) was measured during mitochondrial targeting. Basal OCR was calculated as baseline OCR – oligomycin OCR for each cell line. Spare respiratory capacity (SRC) was calculated as FCCP OCR – baseline OCR for each cell line. N=5 replicates per line. Comparisons made using ANOVA with Dunnett’s multiple comparisons test. ****p<0.0001 relative to MecPK. *p<0.05 relative to KLE. #p<0.05 relative to Hec50. ♮p<0.05 relative to Scr controls.

### 2.5 Mitochondrial defects in MSH2 deficiency-related endometrial cancer confers metabolic vulnerability

To evaluate changes to overall metabolic profile due to mitochondrial defects in MSH2-deficient EC, we performed targeted metabolomics profiling of polar metabolites using ion chromatography mass spectrometry (ICMS) of benign endometrial tissues from control mice (n = 5) and Msh2KO mice (n = 13) as well as EC from Msh2KO mice (n = 7). We compared differentially modulated metabolites in Msh2KO benign versus control benign endometrium and Msh2KO benign versus Msh2KO EC to understand potential metabolic derangements underlying MSH2-deficient carcinogenesis. We also compared Msh2KO EC versus *Pten*^+/-^ EC (N=5) to understand metabolic differences that may be specific to MSH2-deficient carcinogenesis rather than a general carcinogenic phenotype. The statistically significant (FDR<0.05) differences in metabolite abundances between Msh2KO benign (Ben) and control benign are shown in **Table 2**, between Msh2KO benign and Msh2KO benign are shown in **Table 3**, and between Msh2KO EC and *Pten*^+/-^ EC are shown in **Table 4**. Many differentially modulated metabolites in Msh2KO tissues related to carbohydrate metabolism (glycolysis, fermentation, and TCA cycle), nucleotide metabolism, amino acid metabolism, and redox balance, with carbohydrate metabolism and nucleotide metabolism appearing across all three comparisons (21–25).

**Table 2.**
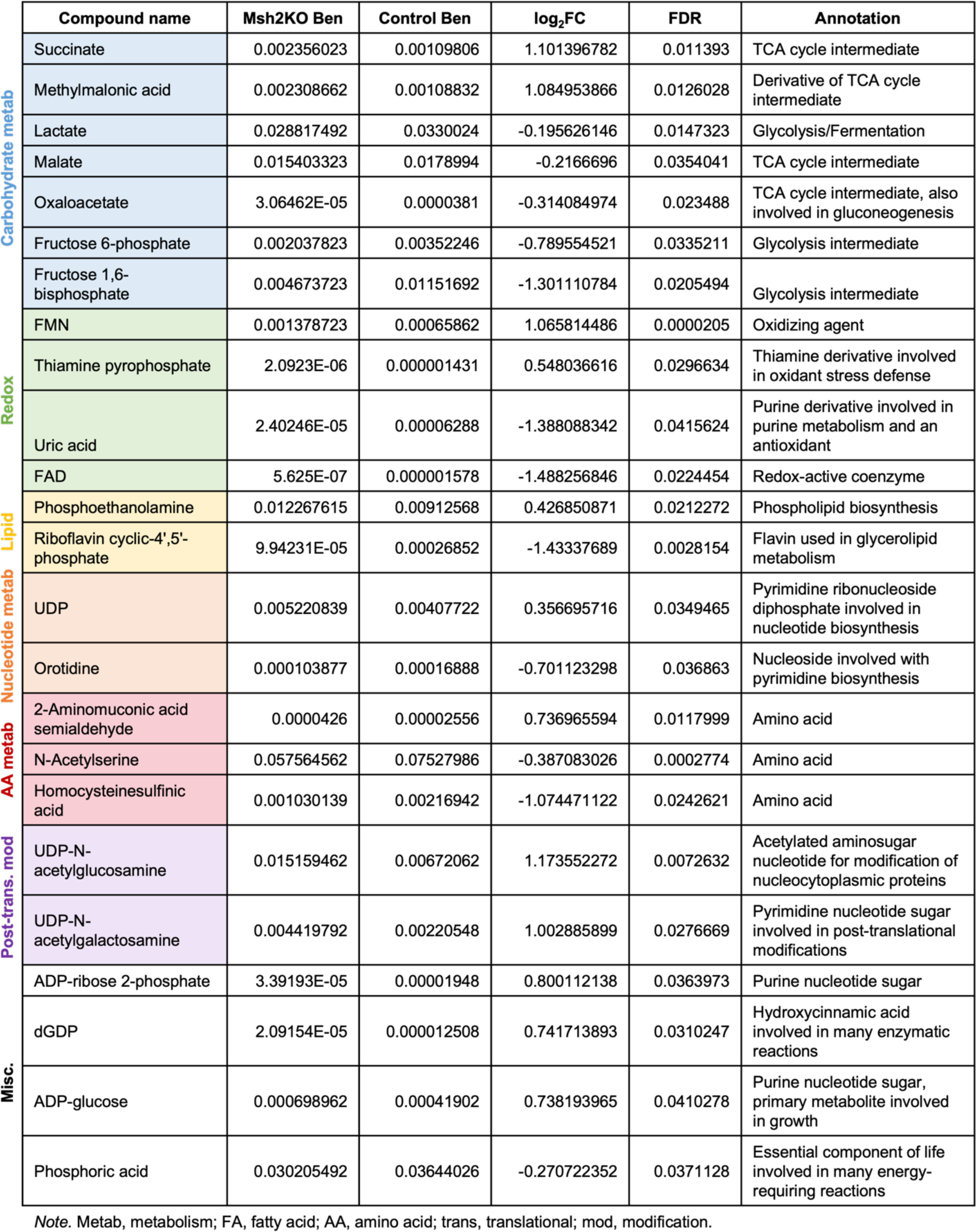
Differentially modulated metabolites in Msh2KO benign relative to control benign endometrium by ICMS polar metabolomics profiling.

**Table 3.**
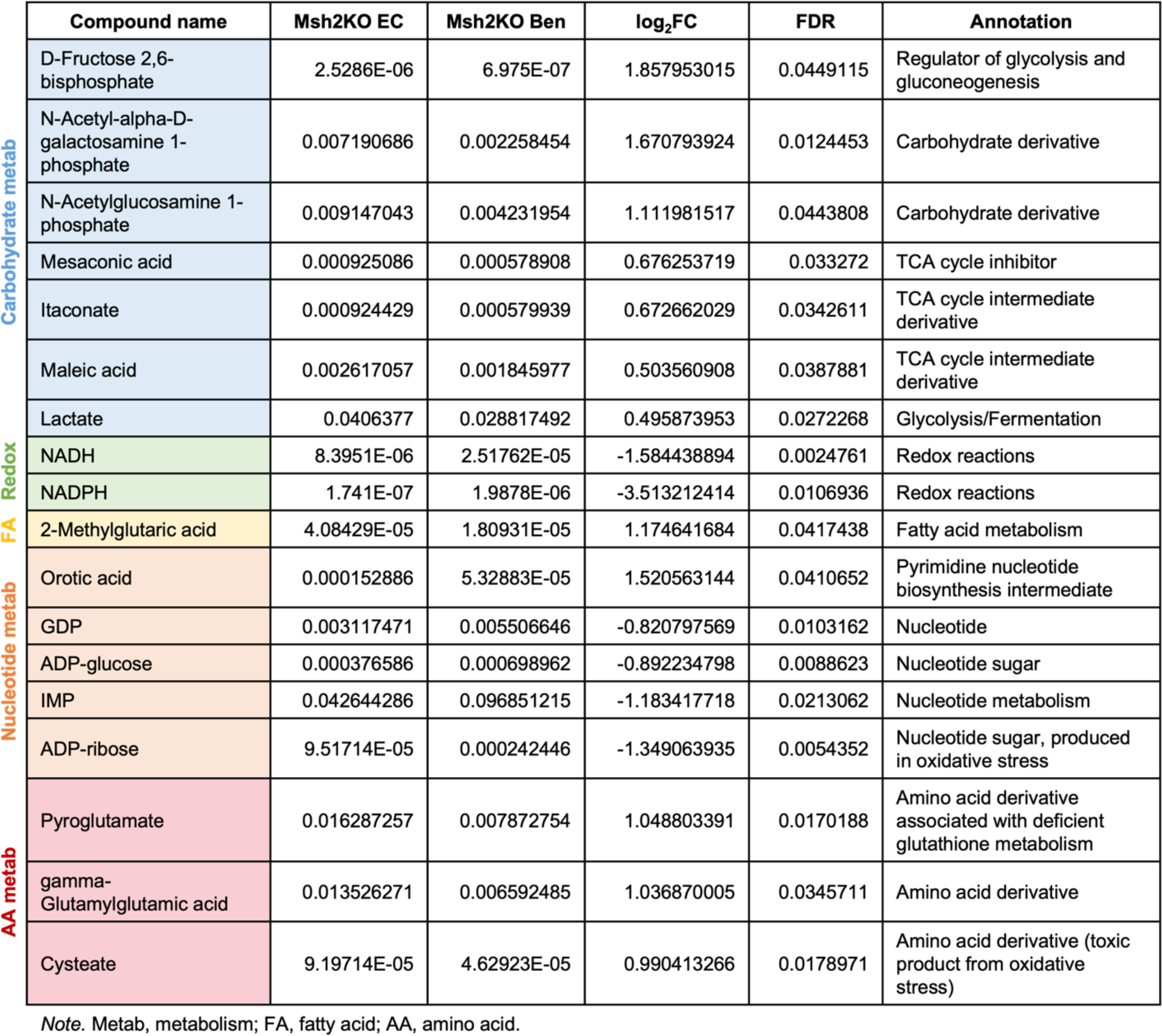
Differentially modulated metabolites in Msh2KO EC relative to Msh2KO benign endometrium by ICMS polar metabolomics profiling.

**Table 4.**
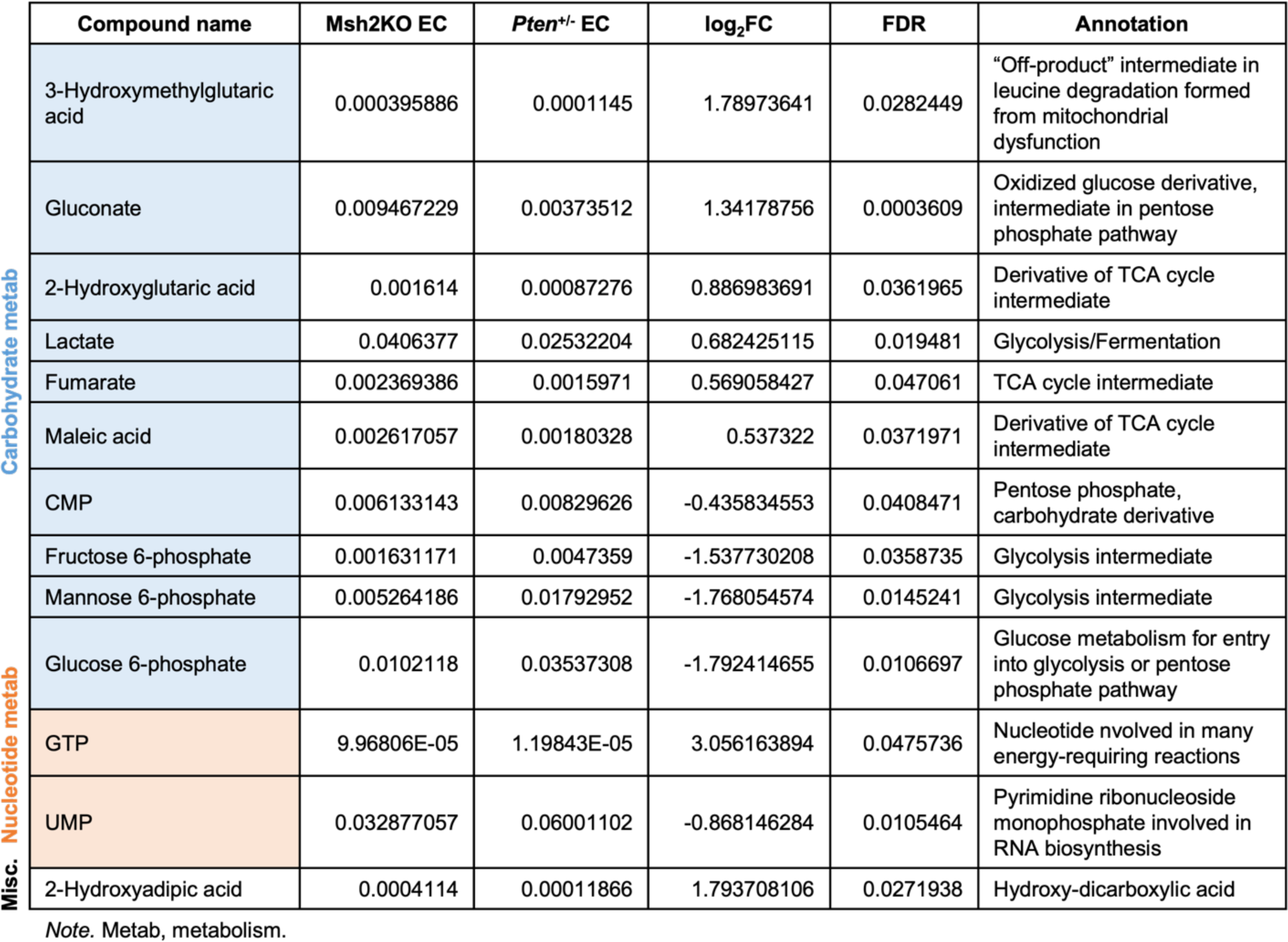
Differentially modulated metabolites in Msh2KO EC relative to *Pten*^+/-^ EC by ICMS polar metabolomics profiling.

Given that our findings suggested mitochondrial dysfunction in MSH2-deficient EC cells thus far, we evaluated further the differences in carbohydrate metabolism as it relates to entry into and exit from the TCA cycle, which takes place in the mitochondrial matrix and depends on intact mitochondrial integrity. Msh2KO tumor tissues (Tum) showed significantly increased abundance of lactate compared to Msh2KO benign endometrium (Ben), suggesting that pyruvate may be converted into lactate rather than entering the TCA cycle in Msh2KO EC more so than in Msh2KO benign endometrium (**Figure 6A-B**). Further, Msh2KO EC exhibited significantly increased abundances of each of the derivatives from TCA cycle intermediates, suggesting premature departure from the TCA cycle rather than completing the cycle (**Figure 6C**).

**Figure 6.**
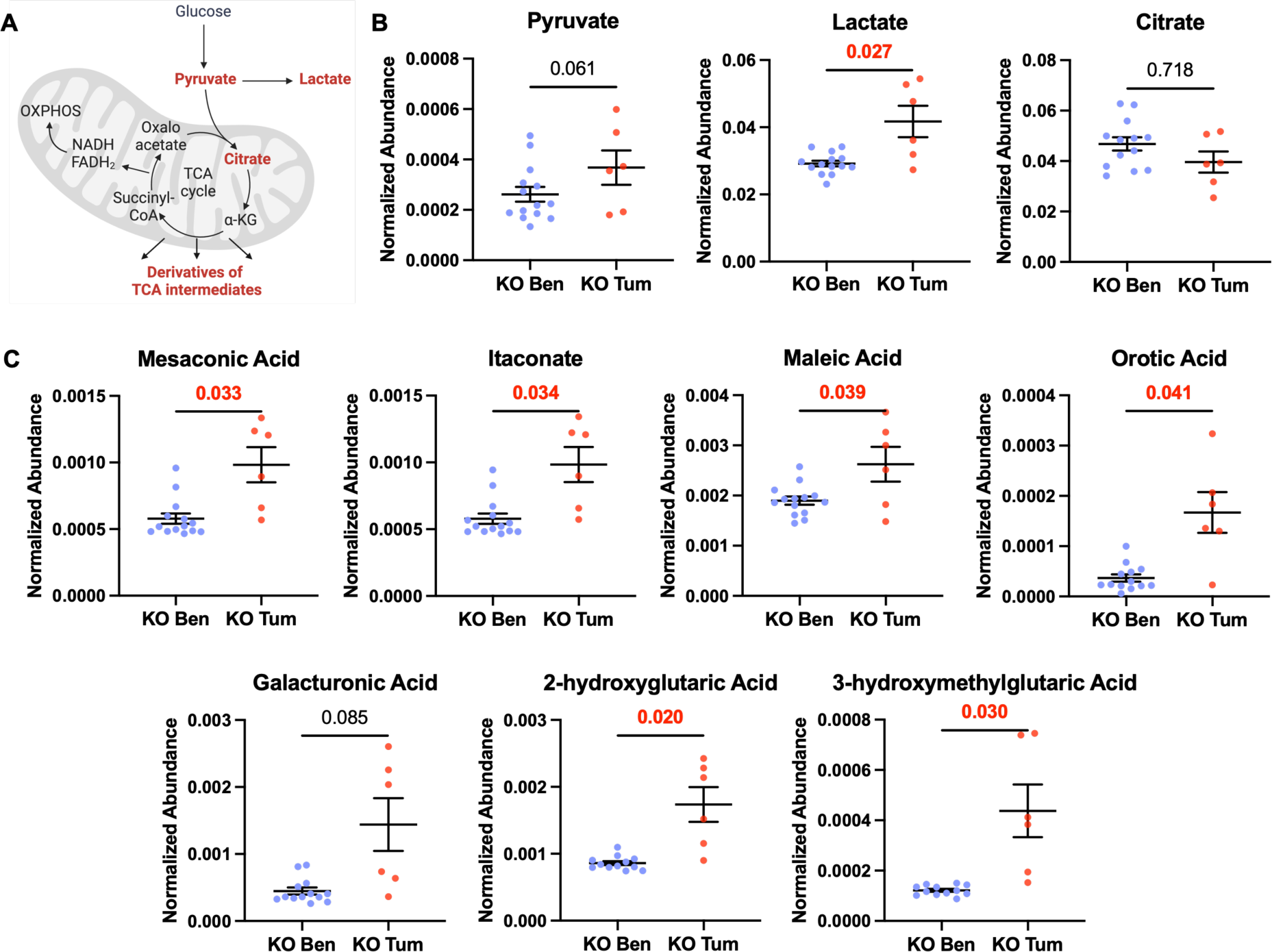
Metabolites indicative of TCA cycle dysfunction were elevated in Msh2KO EC. **(A)** Schematic overview of metabolites related to TCA cycle entry (pyruvate, lactate, citrate) and premature exit from TCA cycle (derivatives of TCA cycle intermediates). **(B)** Quantification of relative peak metabolite abundance of those related to TCA cycle entry (pyruvate, lactate, and citrate) between Msh2KO normal endometrium (blue) and Msh2KO EC (red) tissues using ICMS polar metabolomics profiling. **(C)** Quantification of relative peak metabolite abundance of derivatives of TCA cycle intermediates (mesaconic acid, itaconate, maleic acid, orotic acid, galacturonic acid, 2-hydroxyglutaric acid, 3-hydroxymethylglutaric acid) in Msh2KO normal endometrium (blue) and Msh2KO EC (red) tissues using ICMS polar metabolomics profiling.

Given their mitochondrial dysfunction and evidence of metabolic reprogramming, we determined whether MSH2-deficient EC cells rely on non-mitochondrial pathways for survival. In glucose metabolism, cells can generate ATP through glycolysis followed by fermentation (e.g., producing lactate in anaerobic conditions) or by shuttling pyruvate into the mitochondria for the TCA cycle and oxidative phosphorylation. Overall glycolysis proceeds faster than galactolysis and the surplus NADH generated from glucose preferentially reacts with pyruvate to form lactate. In galactose metabolism, the reliance on glycolysis alone for ATP production is less efficient because galactose metabolism through glycolysis yields less ATP. Thus, cells are more dependent on the TCA cycle and oxidative phosphorylation for sufficient ATP generation. The NADH formed during galactolysis is preferentially used for full oxidation within mitochondria. Therefore, treating cells with galactose-containing media in place of glucose-containing media induces cells to rely more on mitochondria for energy production (26,27). While galactose treatment significantly reduced viability of most mouse and human EC cell lines, MSH2-deficient EC cells had significantly reduced survival after galactose treatment compared to MSH2-intact counterparts, revealing that their reliance on non-mitochondrial pathways for survival represents a metabolic vulnerability (**Figure 7**).

**Figure 7.**
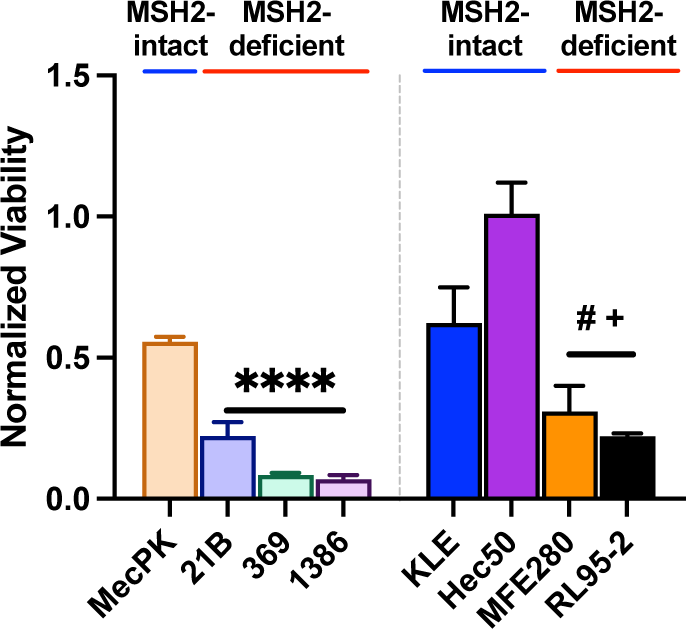
MSH2-deficient EC cells are highly vulnerable to metabolic targeting. MSH2-intact and - deficient murine and human EC cells were treated with glucose-free media supplemented with 4.5 g/L galactose for 72 h. Viability was measured using CellTiter Glo (Promega). Viability relative to glucose-containing media (4.5 g/L) controls are shown. Comparisons made using ANOVA. ****p<0.0001 relative to MecPK. ^#^p<0.05 relative to KLE. ^+^p<0.0001 relative to Hec50.

## 3. Discussion

Pathway analyses from transcriptomic studies of Msh2KO endometrial tissues revealed mitochondrial dysfunction, and we discovered that a broad phenotype of mitochondrial content reduction may underly that finding. Further, mitochondrial staining in human ECs revealed that mitochondrial content depletion is specific to MSH2-deficient ECs and not simply an artifact of carcinogenesis as a whole. Retained mitochondria in MSH2-deficient EC cells have ultrastructural abnormalities in cristae and mitochondrial membranes, reduced mitochondrial membrane potential, and diminished mitochondrial respiration (oxidative phosphorylation) compared to MSH2-intact counterparts. Our studies show a strong association between MSH2-deficient EC and defects in mitochondrial content, integrity, and function. When investigating broader metabolic consequences of mitochondrial defects, differences in metabolite accumulation during EC development in Msh2KO mice suggest that impaired mitochondrial function causes metabolic reprogramming. Indeed, *in vitro* studies suppressing ATP production from glycolysis and fermentation to enhance reliance on oxidative phosphorylation exposed greater vulnerability of MSH2-defiicent EC cells through reduced cell viability compared to MSH2-intact EC cells.

Together, these novel studies indicate that diminished mitochondrial content and damage to retained mitochondria are consequences of MSH2 loss in EC and confer mitochondrial dysfunction, including reduced mitochondrial respiration and increased dependence on non-mitochondrial metabolic pathways. The mechanism for how MSH2 loss causes the mitochondrial phenotype is an area for future study. Ultimately, these findings suggest that mitochondrial dysfunction and metabolic reprogramming could be leveraged as novel biomarkers for EC development and/or targeted for cancer interception.

The observed mitochondrial phenotype as a consequence of MSH2 loss reveals that MMRd has broader effects than solely DNA hypermutability. With such knowledge and characterization of additional carcinogenic pathways in EC, new opportunities for risk stratification, surveillance, and prevention may arise. Future studies elaborating on the timing of mitochondrial defects during EC development and how they provide a selective advantage during carcinogenesis will be paramount to understanding the utility in a clinical context. Nonetheless, this research represents the first known discovery of the association between LS-EC and mitochondrial defects.

To evaluate timing of mitochondrial defects during EC development, measuring mitochondrial content in tissue specimens from endometrial samples throughout the EC development spectrum would be necessary. Further, longitudinal samplign, such as from patients with LS that have surveillance biopsies, would allow for evaluation of mitochondrial content over time with potential progression to pre-cancerous lesions or EC. Lastly, comparisons between benign and/or pre-cancerous tissues from patients with and without LS would aid in determining specificity and timing of mitochondrial aberrations during LS-EC development.

Our study evaluated MSH2-deficient EC to determine additional mechanisms for LS-EC development. However, whether the mitochondrial aberrations are MSH2-specific or relevant to LS more broadly remains to be studied. Additionally, evaluation of MMRd via *MLH1*-deficiency (most commonly via hypermethylation) is an area for future study.

How MSH2-deficiency or MMRd lead to mitochondrial content reduction is an important area for further studies. MSH2 loss and/or MMRd may lead to increased mutations in nuclear-encoded mitochondrial genes. However, we did not observe convergent mutations in nuclear-encoded mitochondrial genes in MSH2-deficient ECs in our study. Another potential mechanism could be hypermethylation of nuclear-encoded mitochondrial genes. Although similar studies are lacking in EC, LS-related colorectal cancers exhibit broad epigenetic alterations (28), lending rationale to this hypothesis. Lastly, questions arise regarding whether MSH2 and other MMR proteins participate in mitochondrial DNA repair as with nuclear DNA repair. Limited studies have evaluated mammalian DNA MMR protein function in the mitochondria, but colocalization studies have found that while MLH1 appears to colocalize with mitochondria and may participate in mitochondrial DNA repair, MSH2 does not (29–31). Sequencing mitochondrial DNA would aid in understanding whether deficient mitochondrial DNA repair may underly our findings. Finally, rather than a direct relationship between MMR proteins and mitochondria, the mitochondrial phenotype could be an indirect consequence of MMRd. Understanding the underlying mechanism whereby MSH2 loss leads to mitochondrial dysfunction will be imperative in determining opportunities for therapeutic targeting in EC prevention and interception.

The mitochondrial phenotype we identified in MSH2-deficient EC may represent a broader signal of stress rather than simply a direct consequence of MSH2 loss. Mitochondrial content reduction occurs in the setting of ongoing cellular stressors, including due to prolonged exposure to elevated glucocorticoids, insulin, and oxidative stress (as reviewed in (33)), which are risk factors in EC development. Perhaps chronic activation of cellular stress pathways in the setting of MSH2 deficiency would contribute to mitochondrial content reduction under similar mechanisms. Such mitochondrial reduction contributes to reduced “resilience” to additional stressors and could contribute to the multiplicative carcinogenic risk of other EC risk factors (such as conditions that increase exposure to unopposed estrogen, obesity, and/or insulin-resistance) in addition to LS (34). Viewing mitochondrial reduction as a phenotype of accumulated “risk” calls for investigation of mitochondrial content changes in other prolonged risk conditions for EC, such as obesity. Excitingly, this would establish a new opportunity for biomarker-based risk stratification in EC based on mitochondrial factors.

In conclusion, this research represents the first descriptions of mitochondrial aberrations due to MSH2 loss in LS-related EC development. These studies reveal that mitochondrial dysfunction is a noncanonical consequence of MMRd in LS-related endometrial carcinogenesis. Biomarkers for mitochondrial defects may be useful in screening or risk stratification purposes for EC. Further, metabolic vulnerability represents a key opportunity that may be leveraged for preventive or therapeutic purposes. Ultimately, these findings add to our understanding of endometrial carcinogenesis and support rationale for future studies on mitochondrial biology in EC.

## 4. Materials and Methods

### 4.1 Generation of PR-Cre Msh2(LoxP/LoxP) mice

Mouse studies were approved by the IACUC at the University of Texas MD Anderson Cancer Center. Progesterone receptor Cre knock-in (PR-Cre) mice were developed previously (19) and were generously gifted by Dr. John Lydon. Msh2(LoxP/LoxP) mice were developed previously (20) and generously shared by Dr. Kucherlapati.

Msh2(LoxP/LoxP) mice were crossed with PR-Cre recombinase mice to generate PR-Cre Msh2(LoxP/+) mice, then intercrossed to create PR-Cre Msh2(LoxP/LoxP) mice (termed Msh2KO). Control mice have Msh2(LoxP/LoxP) gene but lack the PR-Cre recombinase. Offspring from these crosses were genotyped by PCR using previously established methods (19,20) and backcrossed to wildtype C57BL/6J mice. SNP typing was utilized for confirmation of C57BL/6J background using Genetically Engineered Mouse Facility (MD Anderson Cancer Center).

### 4.2 Histologic profiling of mouse tissues

Mice were humanely sacrificed and uterine tissues were harvested, then fixed in 10% formalin and embedded in paraffin. Formalin-fixed paraffin-embedded (FFPE) uterine tissues stained with hematoxylin and eosin (H&E) were examined by a veterinary pathologist for histologic determination.

### 4.3 Human sample studies

Formalin-fixed, paraffin embedded (FFPE) tumors resected from endometrial cancer patients or patients undergoing hysterectomy for other indications were obtained from the University of Texas MD Anderson Cancer Center Gynecologic Oncology Tissue Bank under protocols approved by the Institutional Review Board. Relevant clinical information for human tissues utilized can be found in **Supplementary Table 1**.

### 4.4 Immunohistochemistry

For IHC staining, slides were stained using the Leica Bond Rx Autostainer system. Tissue sections were labeled as single IHC staines for antigens using MSH2 antibody (2017S, Cell Signaling Technology) or TOM20 antibody (42406, Cell Signaling Technology). Slides were dewaxed in Bond Dewax solution (AR9222) and hydrated in Bond Wash solution (AR9590). Heat induced antigen retrieval was performed at 100°C in Bond-Epitope Retrieval solution 1 pH-6.0 (AR9961). After pretreatment, slides were incubated for 1h with MSH2 at 1:300 or TOM20 at 1:200 followed by ready-to-use Novolink Polymer (RE7260-K) secondary. Antibody detection with 3,3’-diaminobenzidine (DAB) and hematoxylin counterstain were performed using the Bond Intense R detection system (DS9263, Leica Biosystems). Stained slides were dehydrated and coverslipped with Cytoseal 60 (23-244256, Fisher Scientific). A positive control and negative control (no primary control) slide was included for this assay. IHC stained slides were digitally imaged in Aperio AT2 (Leica) using 20x objective. For TOM20 quantitation, regions of interest were annotated in Aperio ImageScope software then analyzed using the Cytoplasmic v2 algorithm in the software to determine percent TOM20-high and percent TOM20-low cells with a signal intensity threshold for high expression at 162 (defined as 3+ in the Cytoplasmic v2 algorithm).

### 4.5 Transcriptomic profiling of mouse tissues

Harvested uterine tissue from Msh2KO and control mice were embedded in optimal cutting temperature compound (OCT) and rapidly frozen on dry ice. Tissue sections were fixed in 70% ethanol then microdissected using laser-capture microdissection technology, which enables isolation of regions of interest. Due to microscopic tumor size for some lesions, RNA yield was relatively limited. After RNA isolation, samples were profiled using the Mouse Clariom D assay, a microarray-based broad transcriptome-level expression profiling system that could be performed using low input specimens.

### 4.6 Cell line development and culture information from PR-Cre Msh2(LoxP/LoxP) mice

Two-dimensional cell lines have been established from tumor tissue from Msh2KO mice. These three cell lines (21B, 369, and 1386) were developed from histologically and molecularly distinct tumors and enable mechanistic study of pathways underlying Msh2KO cancer signaling. These are compared with another cell line derived from the MecPK (*Pten*-null, *Kras*-mutant) mouse model for EC which has intact MSH2 expression (35).

Primary mouse cell lines (MecPK, 21B, 369 and 1386) were maintained in DMEM/F12 supplemented with 10% FBS (HyClone), 10 ml/L penicillin/streptomycin solution (Gibco), and 100 mg/ml Primocin (Invivogen). Cell lines were authenticated and tested for mycoplasma using the Characterized Cell Line Core Facility at MD Anderson Cancer Center.

### 4.7 Human cell culture

Human cell lines were purchased from American Type Culture Collection (ATCC) or European Collection of Authenticated Cell Cultures (ECACC). KLE, Hec59, and RL95-2 cells were maintained in DMEM/F12 supplemented with 10% FBS (HyClone), 10 ml/L penicillin/streptomycin solution (Gibco), and 100 mg/ml Normocin (Invivogen). Hec50, AN3CA, and MFE-280 cells were maintained in MEM supplemented with 10% FBS (HyClone), 10 ml/L penicillin/streptomycin solution (Gibco), and 100 mg/ml Normocin (Invivogen). Hec1a and Hec1b cells were maintained in McCoy’s 5a media supplemented with 10% FBS (HyClone), 10 ml/L penicillin/streptomycin solution (Gibco), and 100 mg/ml Normocin (Invivogen).

### 4.8 Immunocytochemistry/Immunofluorescent staining of cells *in vitro*

Cells were seeded onto 0.17 mm (#1.5)-thick glass coverslips (Warner Instruments) pre-coated overnight with 60 μg/ml Collagen I (Gibco). For TOM20 staining, cells were seeded on collagen-coated coverslips and fixed as above. Background Sniper (Biocare Medical) was used to block nonspecific staining, then anti-TOM20 Rabbit mAb (Cell Signaling Technology) was diluted 1:100 in Background Sniper and applied to coverslips overnight at 4°C. Following PBS wash, coverslips were incubated in Goat anti-Rabbit IgG (H+L) highly cross-adsorbed Alexa Fluor Plus 488-conjugated secondary antibody (Invitrogen) at a concentration of 1:1000 in Background Sniper for 1 hour at room temperature. Coverslips were washed, then counterstained with HCS CellMask Deep Red and Hoescht 33342 Solution at concentrations as above for 30 minutes at room temperature. Coverslips were washed and mounted as above.

Imaging of stained cells was conducted with the 63X/1.46 Oil objective on the Zeiss Laser-point Scanning Microscope (LSM) 880 with Airyscan module for ultra-high-resolution imaging in the Advanced Microscopy Core at MD Anderson Cancer Center. Raw .czi files were processed using the Zen software into deconvoluted Airyscan images and analyzed using CellProfiler (Broad Institute). TOM20 intensity was quantified relative to cell area.

### 4.9 Transmission electron microscopy

Cells seeded on tissue culture-treated plates were fixed with 3% glutaraldehyde and 2% paraformaldehyde in 0.1 M cacodylate buffer (pH 7.3), then washed in 0.1 M sodium cacodylate buffer. Processing and imaging took place at the High Resolution Electron Microscopy Core Facility at MD Anderson Cancer Center. Samples were treated with 0.1% Millipore-filtered cacodylate buffered tannic acid, post-fixed with 1% buffered osmium tetroxide, and stained en bloc with 0.1% Millipore-filtered uranyl acetate. The samples underwent serial dehydration in increasing concentrations of ethanol then infiltrated and embedded in LX-112 medium. The samples underwent polymerization in a 60°C oven for 72 h, and ultrathin sections were cut using Leica Ultracut microtome (Leica, Deerfield, IL). Sections were mounted on formvar-coated single slot copper grids and stained with uranyl acetate and lead citrate in a Leica EM Stainer.

The stained samples were examined in a JEM 1010 transmission electron microscope (JEOL USA, Inc., Peabody, MA) using an accelerating voltage of 80 kV. Digital images were obtained using an Advanced Microscopy Techniques imaging system (Advanced Microscopy Techniques Corp, Danvers, MA).

### 4.10 Lentivirus infection for MSH2 knockdown

KLE and Hec50 cells were transduced with lentivirus containing a scrambled shRNA sequence or one of three shRNA sequences complementary to the *MSH2* mRNA. Lentivirus containing U6 promoter-based shRNA vectors were purchased from VectorBuilder (Chicago, IL).

On day 1, cells were trypsinized and resuspended in growth media at a density of 200,000 cells per 1 ml. Polybrene (VectorBuilder) was added to the cell suspension at a volume of 1:1000 and virus was added at a multiplicity of infection (MOI) of 10 transducing units per cell. Each 1 ml suspension containing cells, polybrene, and virus was seeded in one well of a 6-well tissue culture dish. On day 2, virus-containing media was discarded and replaced with fresh growth media. On day 3, growth media was replaced with media containing puromycin (10 μg/ml) for antibiotic selection. Selection was confirmed by visualizing fluorescence of cells infected with an EGFP-expressing control virus.

### 4.11 Mitochondrial membrane potential flow cytometric assay

Live cells were trypsinized, washed, then stained with JC-1 (Invitrogen), a cationic carbocyanide dye that accumulates in mitochondria. JC-1 exists as a monomer at low concentrations and yields green fluorescence. At higher concentrations, JC-1 aggregates form that exhibit red fluorescence. JC-1 accumulation in mitochondria is mitochondrial membrane potential-dependent, so the JC-1 flow assay serves as a sensitive marker for mitochondrial membrane potential. JC-1 staining at a concentration of 10 μg/ml in Live Cell Imaging Solution (Invitrogen) at 37°C for 10 minutes with agitation. Following washing with PBS, cells were analyzed using flow cytometry at the Flow Cytometry and Cellular Imaging Core Facility at MD Anderson Cancer Center. Fl1 and Fl2 fluorescence was measured on the Beckman Coulter Gallios. Ratio of Fl1 to Fl2 represents mitochondrial membrane potential.

### 4.12 Mitochondrial stress tests

Cells were seeded on Seahorse XFe96 cell culture microplates (Agilent). On the day of the assay, cells were equilibrated for 1 h in XF Base Assay Medium supplemented with 10 mM glucose, 1 mM sodium pyruvate, and 2 mM L-glutamine in a CO_2_-free incubator at 37°C. Oligomycin (1.5 μM), carbonyl cyanide-4 (trifluoromethoxy) phenylhydrazone (FCCP) (1 μM), and rotenone A/antimycin (0.5 μM) were loaded into injection ports on the pre-hydrated Seahorse XFe96 sensor cartridge and loaded into the Seahorse XFe96 bioanalyzer for calibration. The microplate was then loaded into the Seahorse XFe96 bioanalyzer for analysis using the XF Cell Mito Stress Test assay template file on Wave software. Parameters of interest are automatically calculated in the Seahorse XF Mito Stress Test Report Generator from Wave data and exported to Excel.

### 4.13 Baseline metabolomics profiling by ion chromatography-mass spectrometry

Flash frozen uterine tissues were homogenized with a Precellys Tissue Homogenizer and metabolites were extracted in ice-cold 0.1% Ammonium hydroxide in 80/20 (v/v) methanol/water. Extracts were centrifuged at 17,000 x *g* for 5 min at 4°C and supernatants were transferred to clean tubes and dried by evaporation under nitrogen. Dried extracts were reconstituted in deionized water. Samples of 5 μl were injected for analysis by IC-MS. IC mobile phase A (MPA; weak) was water, and mobile phase B (MPB; strong) was water containing 100 mM KOH. A Thermo Scientific Dionex ICS-6000+ system included a Thermo IonPac AS11 column (4 μm particle size, 250 x 2 mm) with column compartment kept at 35°C. The autosampler tray was chilled to 4°C. The mobile phase flow rate was 350 μl/min and gradient from 1 mM to 100 mM KOH was used. The total run time was 55 min. To assist the desolvation for better sensitivity, methanol was delivered by an external pump and combined with the eluent via a low dead volume mixing tee. Data were acquired using a Thermo Orbitrap IQ-X Tribrid Mass Spectrometer under ESI negative ionization mode at a resolution of 240,000. Raw data files were imported to Thermo Trace Finder 5.1 software for final analysis.

The peak area (raw relative abundance) of each metabolite was normalized by dividing by the total peak area of the respective sample to generate normalized metabolite relative abundance values.

### 4.14 Galactose experiments *in vitro*

Cells were treated for 72 h with glucose-free medium supplemented with 4.5 g/L galactose (Sigma) or 4.5 g/L glucose (Sigma) as the vehicle control. Cell viability was measured using CellTiter Glo 2.0 (Promega).

### 4.15 Statistical considerations and data analyses

All statistical methods were performed on GraphPad Prism version 7 unless otherwise indicated. Comparison of continuous variables between two groups was performed using Student’s *t*-test. Comparison of continuous variables between greater than two groups was performed using ANOVA with post-hoc Tukey or Dunnett’s test as indicated. Non-parametric alternatives for t-tests (Mann-Whitney U-test) or ANOVA (Kruskal-Wallis test) were used as indicated where data was not expected to follow normal distribution. Benjamini-Hochberg method was employed to correct for multiple hypothesis testing as indicated.

## List of Supplementary Data

**Supplementary Table 1.**
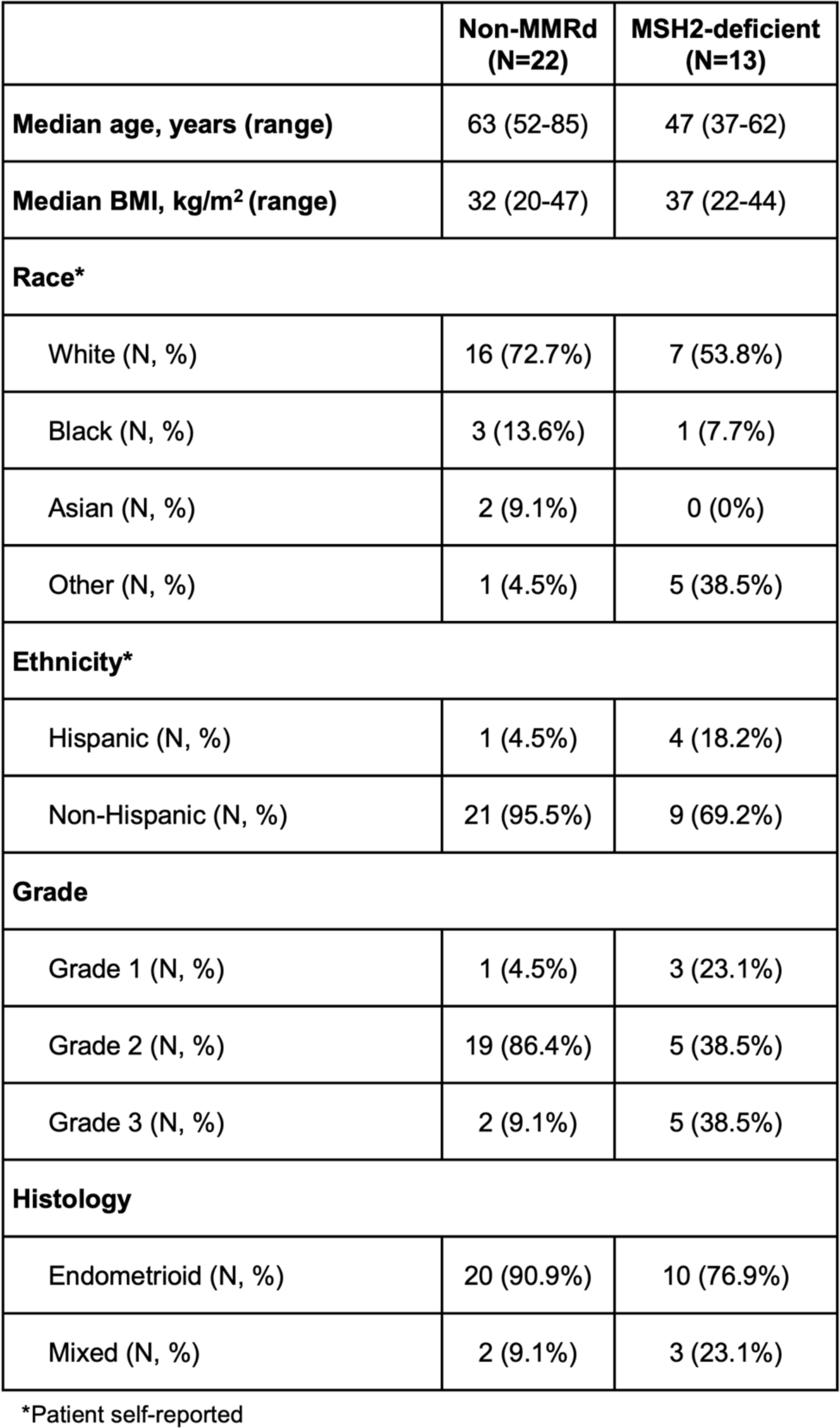
Patient demographic and clinical information for human EC tissues.

|  | **Non-MMRd**<br>**(N=22)** | **MSH2-deficient**<br>**(N=13)** |
| --- | --- | --- |
| **Median age, years (range)** | 63 (52-85) | 47 (37-62) |
| **Median BMI, kg/m<sup>2</sup> (range)** | 32 (20-47) | 37 (22-44) |
| **Race\*** |  |  |
| White (N, %) | 16 (72.7%) | 7 (53.8%) |
| Black (N, %) | 3 (13.6%) | 1 (7.7%) |
| Asian (N, %) | 2 (9.1%) | 0 (0%) |
| Other (N, %) | 1 (4.5%) | 5 (38.5%) |
| **Ethnicity\*** |  |  |
| Hispanic (N, %) | 1 (4.5%) | 4 (18.2%) |
| Non-Hispanic (N, %) | 21 (95.5%) | 9 (69.2%) |
| **Grade** |  |  |
| Grade 1 (N, %) | 1 (4.5%) | 3 (23.1%) |
| Grade 2 (N, %) | 19 (86.4%) | 5 (38.5%) |
| Grade 3 (N, %) | 2 (9.1%) | 5 (38.5%) |
| **Histology** |  |  |
| Endometrioid (N, %) | 20 (90.9%) | 10 (76.9%) |
| Mixed (N, %) | 2 (9.1%) | 3 (23.1%) |

\*Patient self-reported

## Acknowledgments

We gratefully acknowledge the expert technical assistance of Tri Nguyen for tissue processing for histopathology and the MD Anderson Veterinary Medicine team for immunohistochemistry services and expert animal care over these long-term mouse studies. We thank Yongjuan Xia in the Pathology Services Core at the University of North Carolina-Chapel Hill for expert technical assistance with immunohistochemical staining.

